# Reduced *Shmt2* expression impairs mitochondrial folate accumulation and respiration, and leads to uracil accumulation in mouse mitochondrial DNA

**DOI:** 10.1101/2021.04.12.439270

**Authors:** Joanna L. Fiddler, Yuwen Xiu, Jamie E. Blum, Simon G. Lamarre, Whitney N. Phinney, Sally P. Stabler, Margaret E. Brosnan, John T. Brosnan, Anna E. Thalacker-Mercer, Martha S. Field

## Abstract

**Background:** Adequate cellular thymidylate (dTMP) pools are essential for preservation of nuclear and mitochondrial genome stability. Previous studies have indicated that disruption in dTMP synthesis in the nucleus leads to increased uracil misincorporation into DNA affecting genome stability. To date, the effects of impaired mitochondrial dTMP synthesis in non-transformed tissues have been understudied.

**Objective:** This study aimed to determine the effects of decreased serine hydroxymethyltransferase 2 (*Shmt2)* expression and dietary folate deficiency on mitochondrial DNA integrity and mitochondrial function in mouse tissues.

**Methods:** Liver mitochondrial DNA (mtDNA) content, and uracil content in liver mtDNA was measured in *Shmt2^+/-^* and *Shmt2^+/+^* mice weaned onto either a folate-sufficient control diet (2 mg/kg folic acid, C) or a modified diet lacking folic acid (0 mg/kg folic acid, FD) for 7 wks. *Shmt2^+/-^* and *Shmt2^+/+^* mouse embryonic fibroblasts (MEF cells) were cultured in defined culture medium containing either 0 or 25 nM folate to assess proliferative capacity and mitochondrial function.

**Results:** *Shmt2^+/-^* mice exhibited 48-67% reduction in SHMT2 protein levels in tissues. Interestingly, *Shmt2^+/-^* mice consuming the folate-sufficient C diet exhibited a 25% reduction in total folate in liver mitochondria. There was also a >20-fold increase in uracil in liver mtDNA in *Shmt2^+/-^* mice consuming the C diet, and dietary folate deficiency also increased uracil content in mouse liver mtDNA from both *Shmt2^+/+^* and *Shmt2^+/-^* mice. Furthermore, decreased *Shmt2* expression in MEF cells reduced cell proliferation, mitochondrial membrane potential, and oxygen consumption rate.

**Conclusions:** This study demonstrates that *Shmt2* heterozygosity and dietary folate deficiency impair mitochondrial dTMP synthesis, as evidenced by the increased uracil in mtDNA. In addition, *Shmt2* heterozygosity impairs mitochondrial function in MEF cells. These findings suggest that elevated uracil in mtDNA may impair mitochondrial function.

## INTRODUCTION

Folate-mediated one-carbon metabolism (FOCM) is a metabolic pathway that uses folate (vitamin B9) in the form of tetrahydrofolate (THF) cofactors to transfer one-carbon units for biosynthetic reactions. This includes the *de novo* synthesis of purine bases, *de novo* synthesis of thymidylate (dTMP), and the remethylation of homocysteine to methionine (1). Folate metabolism is compartmentalized within the cell, and occurs distinctly within the cytosol, mitochondria, and nucleus (2). The one-carbon moiety from serine provides the primary source of one-carbon units within the cell. Within the mitochondrial compartment, serine is converted to formate. Formate then exits the mitochondria and serves as the primary one-carbon donor for cytosolic and nuclear FOCM (2, 3). FOCM provides one-carbon units for *de novo* purine synthesis and methionine synthesis in the cytosol. Products of mitochondrial FOCM, in addition to formate, include *N-*formylmethionine tRNA and taurinomethylated tRNA for mitochondrial protein synthesis (2,4,5). There is also accumulating evidence that *de novo* dTMP synthesis occurs within all three cellular compartments (cytosol, nucleus, and mitochondria) (2, 6). Disruption of nuclear folate-dependent *de novo* dTMP synthesis results in an elevated deoxyuridine monophosphate/ deoxynucleotide triphosphates (dUTP/dTTP) ratio, which leads to increased uracil misincorporation into DNA (7). Within the nucleus the enzymes required for *de novo* dTMP synthesis form a multi-enzyme complex that is essential to prevent uracil accumulation in nuclear DNA (8) and to maintain genome stability (9).

Most studies investigating the effects of uracil misincorporation into DNA have focused on the nuclear genome. dTMP synthesis in the mitochondria occurs through 1) the salvage pathway via the conversion of thymidine to dTMP by the enzyme thymidine kinase 2 (TK2) and 2) *de novo* synthesis in which the THF-dependent enzymes SHMT2, TYMS, and DHFR/DHFR2 convert dUMP to dTMP (**Figure 1A**) (10). Mitochondrial DNA (mtDNA) replication relies on *de novo* synthesis when nucleotide demand exceeds salvage pathway production. Interestingly, perturbations in dTMP and subsequently dTTP pools for mtDNA replication have previously been associated with mtDNA depletion disorders in humans (11). Furthermore, levels of mitochondrial dNTPs vary in the tissues from young vs. old rats, even in the absence of detectable differences in dNTP levels in whole tissues (12). Compromising the balance of mitochondrial dNTPs affects both the polymerase and exonuclease functions of mitochondrial DNA polymerase, thus increasing errors during DNA replication and compromising mtDNA integrity (13, 14).

**Figure 1.**
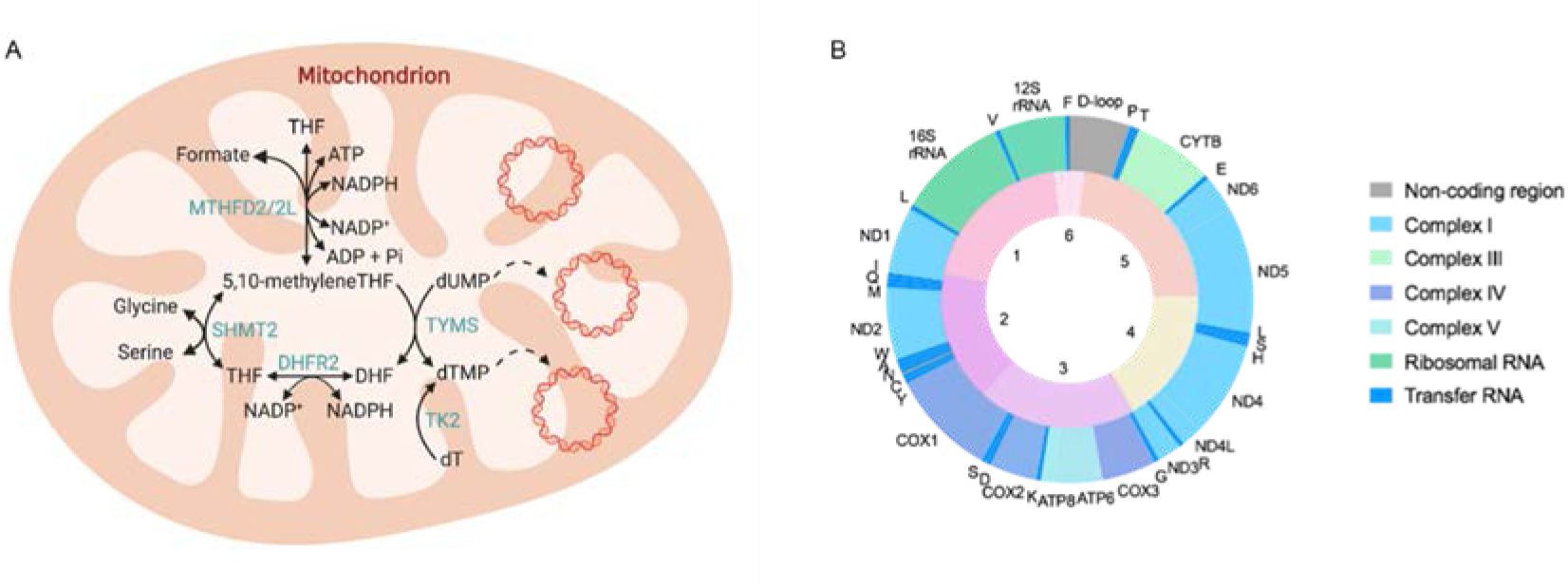
Graphical representations of folate-dependent one-carbon metabolism in the mitochondria and mouse mtDNA genome. A) Folate-dependent one-carbon metabolism in the mitochondrion occurs through the salvage pathway via the conversion of thymidine to dTMP by the enzyme thymidine kinase 2 (TK2) and *de novo* synthesis in which the folate-dependent enzymes SHMT2, TYMS, and DHFR2 convert dUMP to dTMP. B) Schematic of mtDNA with mitochondrial genes reported in the outer ring and mtDNA regions amplified in the real-time PCR assay indicated in the inner ring. DHFR2, dihydrofolate reductase; MTHFD2/2L, methylene THF dehydrogenase; SHMT2, Serine hydroxymethyltransferase; TK2, thymidine kinase; and TYMS, thymidylate synthase.

Maintenance of an adequate cellular dTMP pool is essential to preserve stability of both the nuclear and mitochondrial genomes (15). The only studies quantifying the uracil content in mtDNA in response to impaired mitochondrial FOCM (i.e. decreased SHMT2 levels) or folate deficiency were performed in transformed and/or immortalized cell lines. These studies demonstrate that lack of SHMT2 in Chinese hamster ovary (CHO) cells increases uracil in mtDNA (10) and exposure to folate-deficient culture medium increases uracil in HeLa cell mtDNA (16). In fact, HeLa cell mtDNA is more sensitive to folate deficiency-induced uracil misincorporation than nuclear DNA (16). To date, the effects of *Shmt2* loss and folate deficiency in non-transformed tissues have been understudied. This is important because non-transformed, or primary, tissues often rely more heavily on mitochondria for ATP production than do transformed tissues. Additionally, the physiological consequences of increased uracil in mtDNA remain largely uncharacterized. Therefore, the purpose of this study is to elucidate the metabolic role of SHMT2 in maintaining mtDNA integrity and mitochondrial function across tissues in mice and in a mouse embryonic fibroblast (MEF) cell model.

## EXPERIMENTAL PROCEDURES

### Generation of the Shmt2^+/-^ and Shmt2^-/-^ Mice

*Shmt2* whole-body knockout mice on a C57Bl6/J background were generated by Cyagen Biosciences (Santa Clara, CA). Exon 1 of the mouse *Shmt2* gene (GenBank accession number: NM_028230.4; Ensembl: ENSMUSG00000025403) located on chromosome 10 was selected as the target site. Briefly, the pRP[CRISPR]-hCas9-U6 plasmid was used to generate the targeting vector. The guide RNA (gRNA) sequence (CATCTATACAAGTCCCGGTA) was inserted downstream of the U6 promoter and the hCas9 was inserted downstream of the CBh promotor. Cas9 mRNA and gRNA were *in vitro* transcribed and injected into fertilized mouse embryos. Two chimeric C57Bl6/J male founders (referred to as *Shmt2^+/-^* mice) were obtained and genotyped by PCR using purified tail genomic DNA as a template with the primer pair F: 5’- GGTTTAACATTGGAAGAAACGGAGGA-3’ and R: 5’- GAGTTGTGTACTGGCGGGAGTTGG-3’. The amplicons were purified and DNA sequencing with forward sequencing primer 5’-GTGTTGTAATCCTAAAGCAAGGATTTG-3’ revealed ∼450 bases were missing. The F0 positive heterozygous mice were mated with *Shmt2^+/+^* (wild type) mice to obtain heterozygous F1 generation *Shmt2^+/-^* mice. F1 generation mice were confirmed by genotyping with PCR and DNA sequencing. *Shmt2^+/-^* male and female mice were intercrossed in an attempt to produce *Shmt2^-/-^* mice.

### Animal dietary exposures and tissue collection

All mice were maintained under specific pathogen-free conditions in accordance with standard of use protocols and animal welfare regulations. All study protocols were approved by the Institutional Animal Care and Use Committee of Cornell University. At weaning, male and female *Shmt2^+/-^* and *Shmt2^+/+^* (also referred to as wild-type or WT) offspring (generated from breeding C57Bl6/J female mice and *Shmt2^+/-^* male mice) were randomly assigned to either AIN- 93G control (C) diet (#117814GI; Dyets, Inc., Bethlehem, PA) which contained 2 mg/kg folic acid or to a modified AIN-93G diet lacking folic acid (FD) (#117815GI; Dyets, Inc., Bethlehem, PA). Body weight was measured at intervals of ∼2 weeks for 16 weeks for the growth curve analysis. For all other analyses, 10 week-old mice (7 weeks on defined diets) were used.

Mice were fasted overnight and sacrificed by carbon dioxide asphyxiation followed by cervical dislocation. Blood was collected by cardiac puncture into EDTA treated tubes and plasma was separated by centrifuging at 7,500 rpm for 5 min. Mouse tissues (liver, kidney, brain, skeletal muscle) were washed in ice-cold PBS and snap frozen in liquid nitrogen then stored at - 80 °C or immediately used for liver mitochondrial isolation.

### Mitochondrial isolation

Liver mitochondria was isolated with an OptiPrep (60% iodixanol, Sigma-Aldrich) discontinuous gradient as previously described (16) with slight modifications. Briefly, mouse liver was washed in ice-cold PBS and transferred to 10 mL freshly prepared homogenization medium (0.25 M sucrose, 1 mM EDTA, 20 mM HEPES-NaOH, pH 7.4, and 1:1000 protease inhibitor). Following 20-stroke homogenization with a Dounce homogenizer pestle B, liver was filtered and centrifuged (1,000 x g for 10 min in a fixed-angle rotor) to pellet nuclei. The mitochondrial containing supernatant was transferred and centrifuged (14,000 x g for 12 min at 4 °C) to obtain the crude mitochondrial pellet. The pellets were resuspended in homogenization medium and transferred for 2-3 strokes with a loose-fitting Douce homogenizer pestle. The crude fractions were adjusted to 36% (w/v) iodixanol with 50% iodixanol (diluted in 0.25 M sucrose, 6 mM EDTA, 120 mM HEPES-NaOH, pH 7.4, and 1:1000 fresh protease inhibitor) and transferred to an ultracentrifuge tube. Equal parts of 30% and 10% iodixanol gradient solutions were layered on the mitochondrial fraction and then samples were centrifuged at 50,000 x g for 2 h (Beckman SW41TI). The mitochondrial layer was collected, diluted with 2 mL of homogenization medium, and pelleted by centrifugation (15,000 rpm for 10 min at 4 °C). Pellets were resuspended in 1 mL homogenization medium and centrifuged at maximum speed for 1 min. Final mitochondrial pellets were apportioned for immunoblot assays, microbiological assays, biochemical assays, and mtDNA analysis of uracil concentrations. Mitochondrial fractions were determined to be free of nuclear (lamin A/C) and cytosolic (GAPDH) protein contamination (data not shown).

### Cell culture conditions

Mouse embryonic fibroblasts were isolated at 12.5 days post coitus from C57Bl/6 female mice bred to *Shmt2^+/-^* male mice. Embryos were harvested from the uterus into ice-cold PBS. Heads and viscera were removed and bodies were cut into 1-2 mm pieces and further digested in 0.125% trypsin at 37°C for 20 min. Genotypes were determined for each resulting cell line using PCR as described above. Cells were maintained in alpha-minimal essential medium (alpha-MEM; Hyclone Laboratories) supplemented with 10% FBS and 1% penicillin/streptomycin. For experiments requiring 25 nM and 0 nM folate exposure, cells were cultured in modified alpha-MEM (HyClone) lacking glycine, serine, methionine, folate, and nucleosides which was then supplemented specifically with 10% dialyzed FBS, 200 µM methionine, 1 mg/L pyridoxine, and 0 or 25 nM (6S)-5- formyl-tetrahydrofolate. Cells were monitored for proliferative capacity or allowed to grow for four doublings in modified culture medium prior to harvesting.

### Cell viability

Mouse embryonic fibroblasts were seeded at 1000 cells per well in 96-well plates in 25 nM and 0 nM folate medium with the addition of 0 or 2 mM glycine or formate. The number of total and dead cells was determined at specified time points by co-staining cells with Hoechst 33342 (to identify all cells, Life Technologies) and propidium iodide (to identify dead cells, Thermo Fisher Scientific). Cells were visualized and quantified using a Celigo imaging cytometer (Nexcelom) following the manufacturer’s instructions. The number of live cells was determined by subtracting the number of propidium iodide-positive cells from the Hoechst 33342-positive cells. Data are shown as growth relative to cell number on day 1.

### Mitochondrial oxygen consumption rate and membrane potential

The oxygen consumption rate (OCR) of the MEF cells was measured using the Seahorse XFe24 Extracellular Flux Analyzer (Agilent Technologies). Briefly, cells were cultured in the experimental 25 and 0 nM folate conditions (described above) for 4 doublings, then seeded in the same medium and allowed to adhere for 24 hours or seeded in the same medium with 2 mM glycine or 2 mM formate and allowed to adhere for 48 hours. Prior to the assay, cells were washed twice in Seahorse assay buffer (glucose-free, glutamine-free, pyruvate-free, phenol red-free DMEM (Sigma) supplemented with 1 mM glucose, 2 mM glutamine, and 1 mM pyruvate) and incubated at 37° C for 1 h in a carbon dioxide-free incubator. OCR, extracellular acidification rate (ECAR), or basal respiration were determined following the manufacturer’s instructions for the Cell Mitochondrial Stress Test.

The mitochondrial membrane potential of the MEF cells was determined using JC-1 dye (Cayman) following the manufacturer’s instructions. J-aggregate (excitation/emission = 535/595 nm) and monomer (excitation/emission = 484/535 nm) fluorescence was measured with a SpectraMax M3 (Molecular Devices).

### Immunoblotting

Total protein was extracted following tissue lysis by sonication in lysis buffer (150 mM NaCl, 5 mM EDTA pH8, 1% Triton X-100, 10 mM Tris-Cl, 5 mM dithiothreitol and protease inhibitor) and quantified by the Lowry-Bensadoun assay (17). Proteins were denatured by heating with 6X Laemelli buffer for 5-10 min at 95 °C. Samples were electrophoresed on 8, 10 or12% SDS-PAGE gels for approximately 60-70 min in SDS-PAGE running buffer and then transferred to an Immobilon-P polyvinylidene difluoride membrane (Millipore Corp.) using a MiniTransblot apparatus (Bio-Rad). Membranes were blocked in 5% (w/v) nonfat dairy milk in 1X phosphate-buffered saline (PBS) containing 0.1% Tween-20 for 1 h at room temperature. The membranes were incubated overnight in the primary antibody at 4 °C and then washed with 1X TBS containing 0.1% Tween-20 and incubated with the appropriate horseradish peroxidase – conjugated secondary antibody at 4 °C for 1 h. The membranes were visualized with Clarity and Clarity Max ECL Western Blotting Substrates (Bio-Rad). Antibodies against SHMT2 (Cell Signaling, 1:1000), VDAC (Cell Signaling, 1:1000), and GAPDH (Cell Signaling, 1:2000) were used. For antibody detection, a goat anti-rabbit IgG-horseradish peroxidase-conjugated secondary (Pierce) was used at a 1:15000 dilution. Densitometry was performed with ImageJ (version 1.53a) using VDAC or GAPDH as the control.

### Mitochondrial SHMT2 enzyme activity

Mitochondria were isolated as described above and lysed in 20 mM sodium phosphate buffer, pH 7.2, 10 mM 2-mercaptoethanol, 0.5% Triton X-100. Mitochondrial SHMT2 activity was measured as previously described (18) with slight modifications. Briefly, mitochondrial SHMT2 activity was measured by diluting 20 μl of the lysed mitochondrial fraction to 150 μl with 10 mM potassium phosphate buffer, pH 7.5, 10 mM 2-mercaptoethanol, and 4 nmol tritiated glycine (American Radiochemicals) with both protons of the 2-carbon tritiated. The reaction was initiated by the addition of 4 nmol of (6S)-THF and incubated at 37°C for 4 hours with gentle agitation. Control reactions were performed to correct for background exchange by the addition of 4 nmol of (6S)5-formyl in lieu of (6S)THF. The reaction was terminated by the addition of 1 mL of 50 mM HCl, and the solution was passed through a column containing 0.6 ml of Dowex 50 AG (Bio-Rad) to remove radiolabeled glycine. The column was washed with an additional 1 mL of 50 mM HCl, and the tritiated water was collected and quantified by scintillation counting. Control reactions contained pooled samples for each group and subtracted as the background. Protein concentrations were determined by BCA assay according to the manufacturer’s instructions (Thermo Fisher Scientific).

### Folate concentration analysis

Folate concentration of liver, plasma, mitochondrial fractions purified from liver, and MEF cells was quantified using the *Lactobacillus casei* microbiological assay as previously described (19). Total folates were normalized to protein concentrations for each sample (17).

### Plasma metabolite profile and formate analysis

Total homocysteine, cystathionine, total cysteine, methionine, glycine, serine, α- aminobutyric acid, N,N-dimethylglycine, and N-methylglycine were assayed in mouse plasma by stable isotope dilution capillary GC-MS as described previously (20, 21).

Formate levels were assayed and quantified as previously described (22) with slight modifications. Briefly, the GC-MS system composed of an Agilent gas chromatograph (model 7890B) interfaced with a single quadrupole mass selective detector (MSD 5977B) was used for the analysis with ^18^O2-sodium-formate serving as an internal standard. Peak detection and integration were performed using MassHunter (Version B07.01 SP2, Agilent).

### Mitochondrial DNA content and mass

Total genomic DNA was isolated with the Roche High Pure PCR Template Preparation Kit per manufacturers’ protocol and eluted in DNase/RNase free water. Mitochondrial DNA copy number was quantified by real-time quantitative PCR (Roche LightCycler® 480) as previously described (23), using LightCycler® 480 SYBR Green I Master (Roche) and 15 ng DNA per reaction. Oligonucleotide primers for Mito (F 5’- CTAGAAACCCCGAAACCAAA and R 5’- CCAGCTATCACCAAGCTCGT, B2M (F 5’- ATGGGAAGCCCGAACATACTG and R 5’- CAGTCTCAGTGGGGGTGAAT), and tRNA (F 5’ – 2 CACCCAAGAACA and R 5’ - GGCCATGGGTATGTTGTTA) were purchased from IDT. Mitochondrial mass was determined with the Citrate Synthase Activity Assay Kit (Sigma) according to the manufacturer’s instructions and normalized to the total protein concentration.

### RNA Isolation and cDNA Synthesis

Total RNA was isolated from mouse tissues using RNeasy Mini Kit (Qiagen) or RNA STAT-60 (Tel-test, Inc.) according to manufacturer’s instructions. After isolation, RNA concentration was determined using Nanodrop spectrophotometer (Thermo Fisher Scientific) and relative purity of total RNA was assessed by A_260/280_ ratio. Integrity of RNA was assessed by examining 18S and 28S rRNA by agarose gel electrophoresis. RNA was treated with DNase I (Roche) and then reverse-transcribed with High Capacity cDNA Reverse Transcription Kit (Applied Biosystems).

### Quantitative RT-qPCR and Data Analysis

Relative mRNA expression was determined by RT-qPCR using LightCycler® 480 SYBR Green I Master Mix on a LightCycler® 480 instrument (Roche). All reactions were performed in 10 or 20 µL volumes, including 20 or 40 ng of cDNA and 0.5 µM of the oligonucleotide primers specific to the mRNA of interest. Amplification was performed with a 2 min activation step at 50 °C, 10 min denaturation step at 95 °C, followed by 40 cycles of 95 °C for 15 sec and 60 °C for 1 min. A dissociation curve analysis was performed using the default settings of the software to confirm the specificity of the PCR products. For each target gene, the relative standard curve method was used to analyze data [39]. Oligonucleotide primers for *Shmt2, β-actin,* and *18S* were obtained from Qiagen.

### Quantitative analysis of liver uracil concentrations

Uracil present in liver genomic DNA was determined as previously described (8) with minor modifications. Genomic DNA was isolated using the Roche High Pure PCR Template Preparation Kit per manufacturers’ protocol and eluted in DNase/RNase free water. Liver DNA was treated with RNase A (Sigma-Aldrich), incubated at 37 °C for 15 min, and then purified using a Qiagen PCR purification kit according to the manufacturer’s instructions. DNA concentration was quantified using a Qubit Fluorometric Quantification (Thermo Scientific) and 2 µg DNA was treated with uracil DNA glycosylase (New England Biolabs, Inc.) at 37 °C for 60 min. Following incubation, 50 pg of internal standard [^15^N_6_] uracil (Cambridge Isotope Laboratories, Inc.) was added to each, and the samples and standards were dried in a desiccator for 3 days. Samples were derivitized using 3,5-bis(trifluoromethyl)benzyl bromide and prepared for analysis using Q3 selected ion monitoring on a Shimadzu TQ8030 with gas chromatography and mass spectrum settings as previously described (8) with the addition of correction using the internal standard.

### Quantification of uracil in mtDNA by real-time PCR

Mitochondria were isolated as described above, followed by mtDNA isolation with QIAprep Spin Miniprep Kit (Qiagen). To examine the total uracil content in mtDNA, we designed six primer sets producing amplicons of 646-3,751 bp to cover the entire mtDNA (**Table 1** and Figure 1B). The primers were designed using NCBI/primer-BLAST and synthesized by Integrated DNA technologies (IDT) with HPLC purification. All primer sets yielded a single product, confirming the specificity of the long-run real-time PCR (Figure S4). The reaction mixture was prepared as previously described (24). Serial dilutions of mtDNA were applied as templates for Phusion DNA polymerase (wild-type) and Phusion U DNA polymerase (point mutant) in real-time PCR. While the wild-type polymerase stalls if uracil is detected in the DNA template, the mutant polymerase does not recognize uracil and allows replication of any DNA templates. Quantitative real-time PCR reactions were carried out in a LightCycler 480 II system (Roche) in 384-well plates. Reaction conditions were: pre-incubation at 95 °C for 5 min (1 cycle); denaturation at 95 °C for 15 s, annealing at 59 °C for 20 s and extension at 72 °C for 50 s in the case of region 6 and 3 min in the case of region 1-5 (40 cycles); melting at 95 °C for 5 s, 57 °C for 20 s and 95 °C continuous (1 cycle) and cooling at 40 °C for 30 s. Specific PCR products were confirmed on agarose gels. Uracil content was calculated as previously described (24). Briefly, the *Cq* shift between uracil-containing and reference DNA, when amplified with the two polymerases, was calculated to quantify the amount of uracil in each PCR region in the mtDNA samples. MtDNA from a single *Shmt2^+/+^* mouse consuming the control diet was used as the reference DNA for all samples.

**Table 1.**
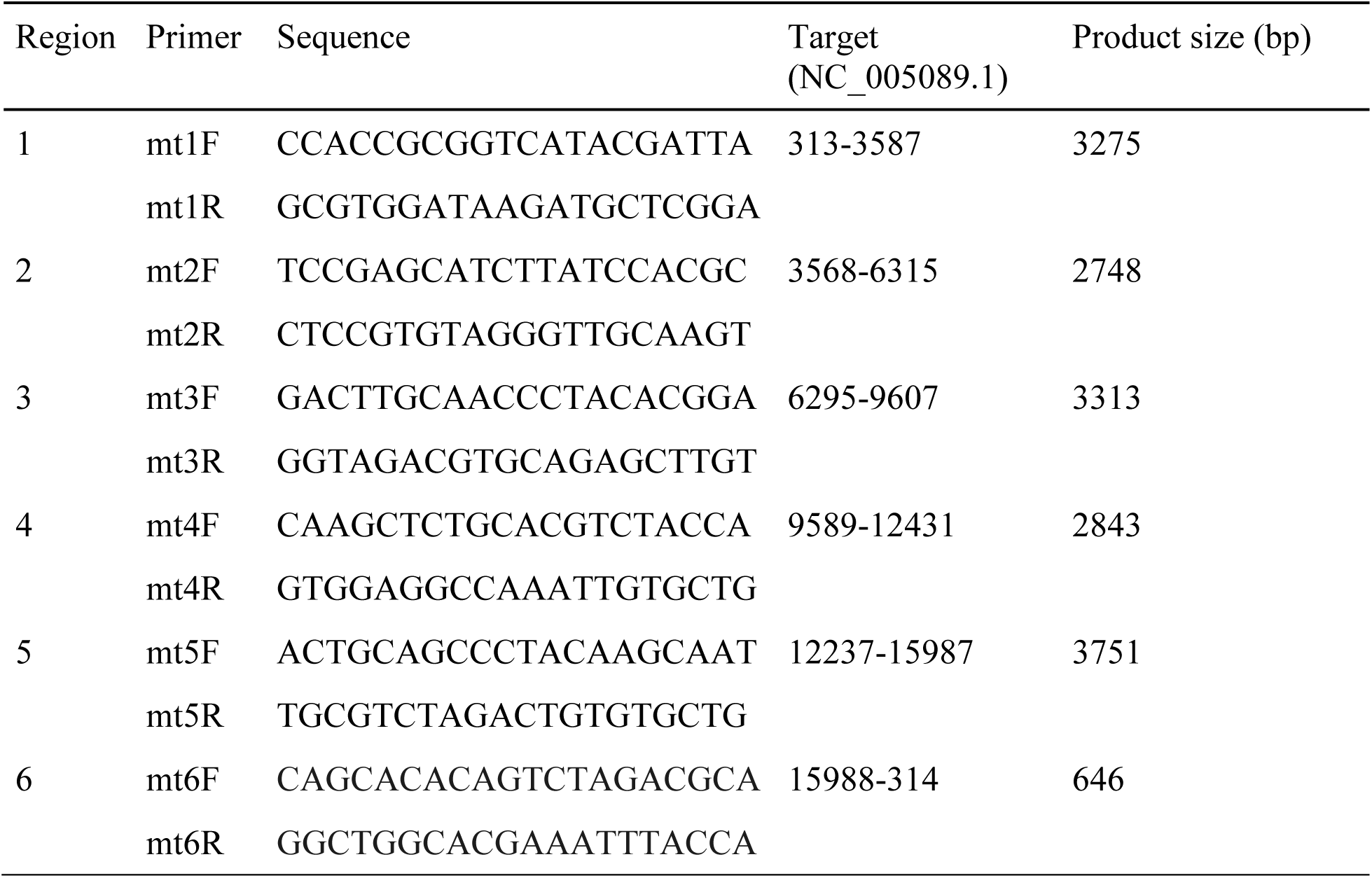
Primers for quantification of uracil in mouse mtDNA.

### In vitro base excision repair of mtDNA for validation of uracil in mtDNA assay

MtDNA was treated with 1 μl uracil-DNA glycosylase (UDG) (NEB, M0280), 1.5 μl Endonuclease IV (NEB, M0304), 0.8 μl *Bst* DNA polymerase, full length (NEB, M0328), 1.7 μl Taq ligase (NEB, M0208), 1 μl NAD^+^ (NEB, B9007) and 1 μl dNTP (Promega, U1515) in NEBuffer 3 in a 50 μl reaction at 37°C for 60 min (25). DNA was then purified by ethanol precipitation. Samples with (uracil-free DNA) and without (uracil-containing DNA) base excision repair were assayed as described above.

### Statistical analyses

JMP® Pro statistical software version 14 (SAS Institute Inc.) was used for all statistical analyses. A growth curve analysis was performed using linear mixed models. In this analysis the model included fixed effects of genotype, diet, and the interaction between diet and genotype. The model allowed for separate intercepts, slopes, and quadratic terms of time for each genotype by diet group and random intercepts, slopes, and quadratic effects at the mouse level. A residual analysis was performed to check the model assumptions of normality and homogeneous variance. F tests using the Kenward-Roger degrees of freedom correction were used to test the statistical significance of the model fixed effects. *p*-values ≤ 0.05 were considered statistically significant. Differences in genotype distribution were analyzed by Chi-square test. For analyses in which *Shmt2^+/+^* and *Shmt2^+/-^* were placed onto either of the two diets (control or folate deficient) results were analyzed by two-way ANOVA followed by Tukey post-hoc analysis. Linear mixed effects models with random effect of embryo, main effects of media, genotype, and time (with time as a continuous variable), and all 2-, 3-, and 4-way interactions were used to determine MEF cell proliferation. Two-way ANOVA with Tukey’s post-hoc analysis was used to determine medium by genotype interaction and main effects of medium and genotype for all other MEF cell analyses. All tests were performed at the 95% confidence level (α = 0.05) and groups were considered significantly different when *p ≤ 0.05*. Descriptive statistics were calculated on all variables to include means and standard deviations.

## RESULTS

### *Shmt2* homozygous knockout is embryonically lethal

*Shmt2* knockout mice were generated on a C57BL/6J background by CRISPR/Cas9- mediated genome engineering. To determine the viability of *Shmt2* knockout mice, the genotype distribution was determined from *Shmt2^+/-^* intercrosses. Among the 47 surviving pups, 38% (n=18) of the pups were *Shmt2^+/+^* (wild-type), 62% (n=29) of the pups were *Shmt2^+/-^* (heterozygous) mice, and no *Shmt2^-/-^* pups were observed (**Table 2**). The observed and expected genotype distributions were compared using a Chi-Square analysis, which indicated embryonic lethality of the homozygous knockout of *Shmt2,* and that *Shmt2* is an essential gene in C57Bl/6J mice (Table 2) as has been reported in a similar model (26).

**Table 2.** Shmt2 mouse viability. Shmt2^+/-^ mice were intercrossed and the progeny genotyped. The expected genotype distribution was calculated by Mendelian distribution and differences in expected and observed genotypes were analyzed by Chi square analysis. *P* values ≤ 0.05 were considered significantly different.

| Genotype | Expected distribution | Observed distribution |
| --- | --- | --- |
| *Shmt2*<sup>+/+</sup> | 11.8 | 18 |
| *Shmt2*<sup>+/-</sup> | 23.5 | 29 |
| *Shmt2*<sup>-/-</sup> | 11.8 | 0 |
| Litters observed |  | 6 |
| Mean litter size, including resorptions (mean ± SD) |  | 7.8 ± 3.9 |
| *p* value, observed versus expected genotype distribution (1:2:1) |  | <0.001 |

### Characterization of *Shmt2^+/-^* mice

*Shmt2^+/+^* and *Shmt2^+/-^* male and female mice were generated by intercrossing C57BL/6J dams to *Shmt2^+/-^* male mice. *Shmt2^+/+^* and *Shmt2^+/-^* mice were weaned at 21 days and randomly assigned to a control (C) or folate-deficient (FD) diet (described in Materials and Methods). The growth curve indicated that linear and quadratic effects of time were significant (*p* < 0.0001 and *p* < 0.0001, respectively), but there was no significant difference in the intercepts, slopes, or quadratics between genotype or diet group groups (*p* > 0.1; **Figure S1**), indicating that neither genotype nor diet affected animal growth. Additionally, the mice did not exhibit any apparent behavioral or pathological abnormalities. After 7 weeks consuming the FD diet, mice had significantly lower folate levels in plasma (85%, *p* < 0.0001), liver (36%, *p* < 0.001) and liver mitochondria (46%, *p* < 0.0001) compared to folate levels in the mice consuming the C diet (**Table 3**). There was a significant diet-by-genotype interaction in plasma folate levels (*p* < 0.05). Interestingly, *Shmt2^+/-^* mice exhibited decreased liver mitochondrial folate levels (*p* < 0.01), even when mice consumed the C diet (Table 3), suggesting that decreased mitochondrial SHMT2 levels impair mitochondrial folate accumulation in the presence of adequate dietary folate.

**Table 3.** Plasma, liver and mitochondrial folate in *Shmt2* null mice. Folate concentrations were determined using the *Lactobacillus casei* assay. Two-way ANOVA with Tukey’s post-hoc analysis was used to determine diet-by-genotype interaction and main effects of diet and genotype with a statistical significance at *p* ≤ 0.05. Data represent means ± SD values. n = 4 per group for plasma and n = 5-6 per group for liver and liver mitochondria. Values not connected by the same letter are statistically different.

| Diet | *Shmt2*<br>genotype | Plasma<br>*ng/ml* | Liver<br>*fmol/μg protein* | Liver mitochondria<br>*fmol/μg protein* |
| --- | --- | --- | --- | --- |
| AIN-93G | *Shmt2*<sup>+/+</sup> | 79.5 ± 4.7<sup>a</sup> | 94.8 ± 15.1<sup>a</sup> | 486 ± 48.5<sup>a</sup> |
| | *Shmt2*<sup>+/-</sup> | 68.0 ± 3.3<sup>a</sup> | 89.8 ± 25.4<sup>a</sup> | 362 ± 108<sup>ab</sup> |
| AIN-93G minus folate | *Shmt2*<sup>+/+</sup> | 9.3 ± 8.4<sup>b</sup> | 65.2 ± 15.9<sup>ab</sup> | 269 ± 64.1<sup>bc</sup> |
| | *Shmt2*<sup>+/-</sup> | 12.4 ± 4.3<sup>b</sup> | 52.4 ± 14.9<sup>b</sup> | 194 ± 39.1<sup>c</sup> |
| *p* value, diet effect |  | <0.0001 | 0.0004 | <0.0001 |
| *p* value, genotype effect |  | 0.155 | 0.267 | 0.006 |
| *p* value, diet by genotype |  | 0.021 | 0.626 | 0.446 |

*Shmt2* gene expression was determined by quantitative reverse transcription PCR in the brain, heart, kidney, liver, soleus and tibialis anterior muscles of the *Shmt2^+/+^* and *Shmt2^+/-^* mice. The *Shmt2^+/-^* mice exhibited markedly reduced *Shmt2* gene expression in the brain (*p* < 0.001), heart (*p* < 0.05), kidney (*p* < 0.05), and liver (*p* < 0.01) compared to the *Shmt2^+/+^* group (**Figure 2A-D**). *Shmt2* gene expression levels appear lower in *Shmt2^+/-^* soleus compared to the *Shmt2^+/+^* mice, however they did not reach statistical significance due to high variability between animals (Figure 2E). Mice fed a FD diet also exhibited reduced *Shmt2* gene expression in the liver (*p* < 0.01) compared to the C group (Figure 2D). Additionally, there was a diet by genotype interaction in *Shmt2* gene expression levels in the tibialis anterior (*p* < 0.01, Figure 2F).

**Figure 2.**
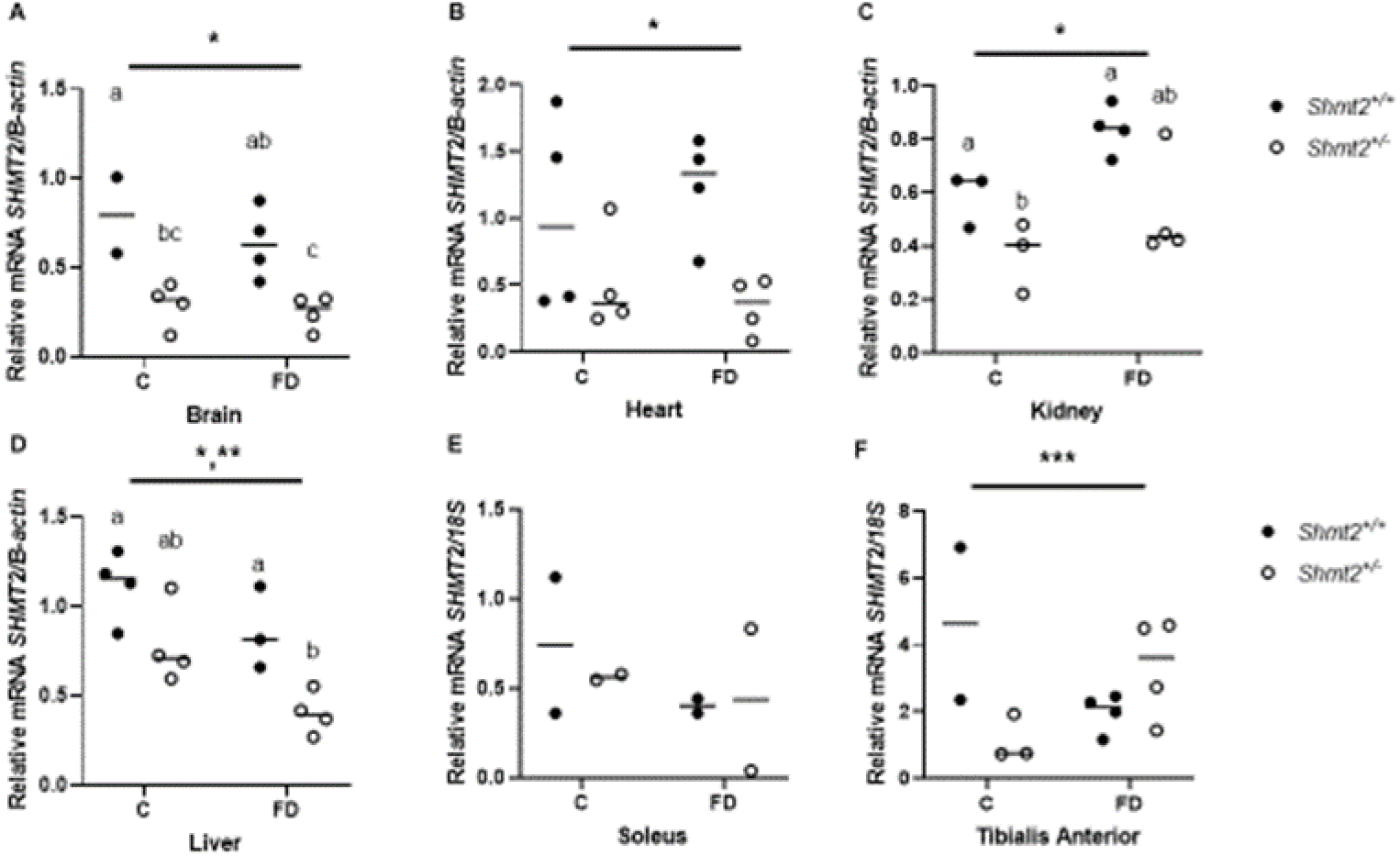
Shmt2 mRNA expression. *Shmt2* mRNA expression in *Shmt2^+/+^* and *Shmt2^+/-^* female mice consuming the C or FD diet for 7 weeks in A) brain, B) heart, C) kidney, D) liver, E) soleus, and F) tibialis anterior. Data are normalized to *β-actin* or *18S* and represent means ± SD. Two-way ANOVA with Tukey’s post- hoc analysis was used to determine diet by genotype interaction and main effects of diet and genotype with a statistical significance at *p* ≤ 0.05. Statistical significant was determined for genotype (*), diet (**), and diet by genotype interaction (***). Levels not connected by the same letter are significantly different, n = 2-3 per group. C, control diet; FD, folate-deficient diet; SHMT2, serine hydroxymethyltransferase 2.

Levels of processed, mitochondrial SHMT2 protein were determined in whole liver, mitochondria purified from liver, brain, heart, kidney, soleus, and tibialis anterior by western blot analyses. In whole liver, *Shmt2^+/-^* mice exhibited 67% lower SHMT2 protein levels than the *Shmt2^+/+^* mice (*p* < 0.01, **Figure 3A**). In liver mitochondria (Figure 3B-C), *Shmt2^+/-^* mice exhibited 58% (*p* < 0.05) lower SHMT2 protein levels and 48% (*p* < 0.05) lower SHMT2 enzyme activity. In brain, (**Figure 4A**), heart (Figure 4B), kidney (Figure 4C), and soleus (Figure 4D), SHMT2 protein levels in *Shmt2^+/-^* mice were 55% (*p* < 0.05), 63% (*p* < 0.01), 79% (*p* < 0.05), and 60% (*p* < 0.001) lower than *Shmt2^+/+^* mice, respectively. SHMT2 protein levels appear lower in *Shmt2^+/-^* tibialis anterior compared to the *Shmt2^+/+^* mice, however they did not reach statistical significance due to high variability between animals (Figure 4E). Interestingly, mice consuming the FD diet exhibited lower SHMT2 protein levels in soleus (*p* < 0.05, Figure 4D).

**Figure 3.**
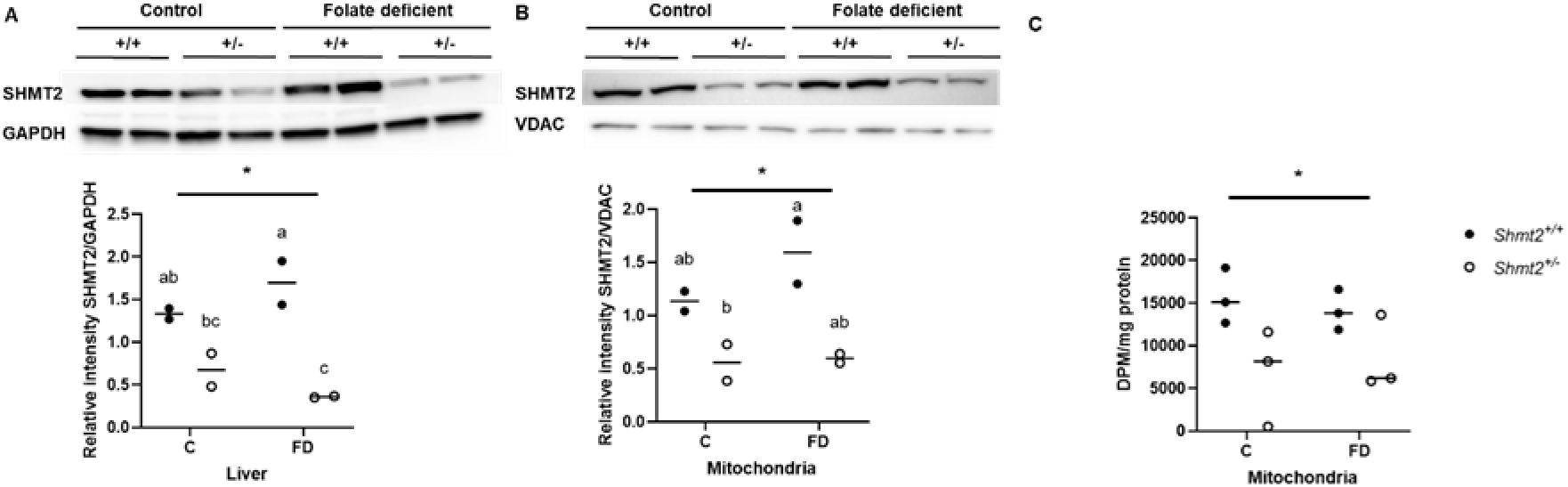
SHMT2 western blot analyses and enzyme activity assay. SHMT2 protein levels and enzyme activity in *Shmt2^+/+^* and *Shmt2^+/-^* female mice consuming the C or FD diet for 7 weeks in A) whole liver protein, B) liver mitochondria protein, and C) enzyme activity in liver mitochondria. Data from female mice are normalized to GAPDH or VDAC. Densitometry was performed using ImageJ. Two-way ANOVA with Tukey’s post-hoc analysis was used to determine diet by genotype interaction and main effects of diet and genotype with a statistical significance at *p* ≤ 0.05. Statistical significant was determined for genotype (*), diet (**), and diet by genotype interaction (***). Levels not connected by the same letter are significantly different, n = 2 per group. C, control diet; FD, folate-deficient diet. GAPDH, glyceraldegyde-3 phosphate dehydrogenase; SHMT2, serine hydroxymethyltransferase 2; VDAC, voltage-dependent anion channel.

**Figure 4.**
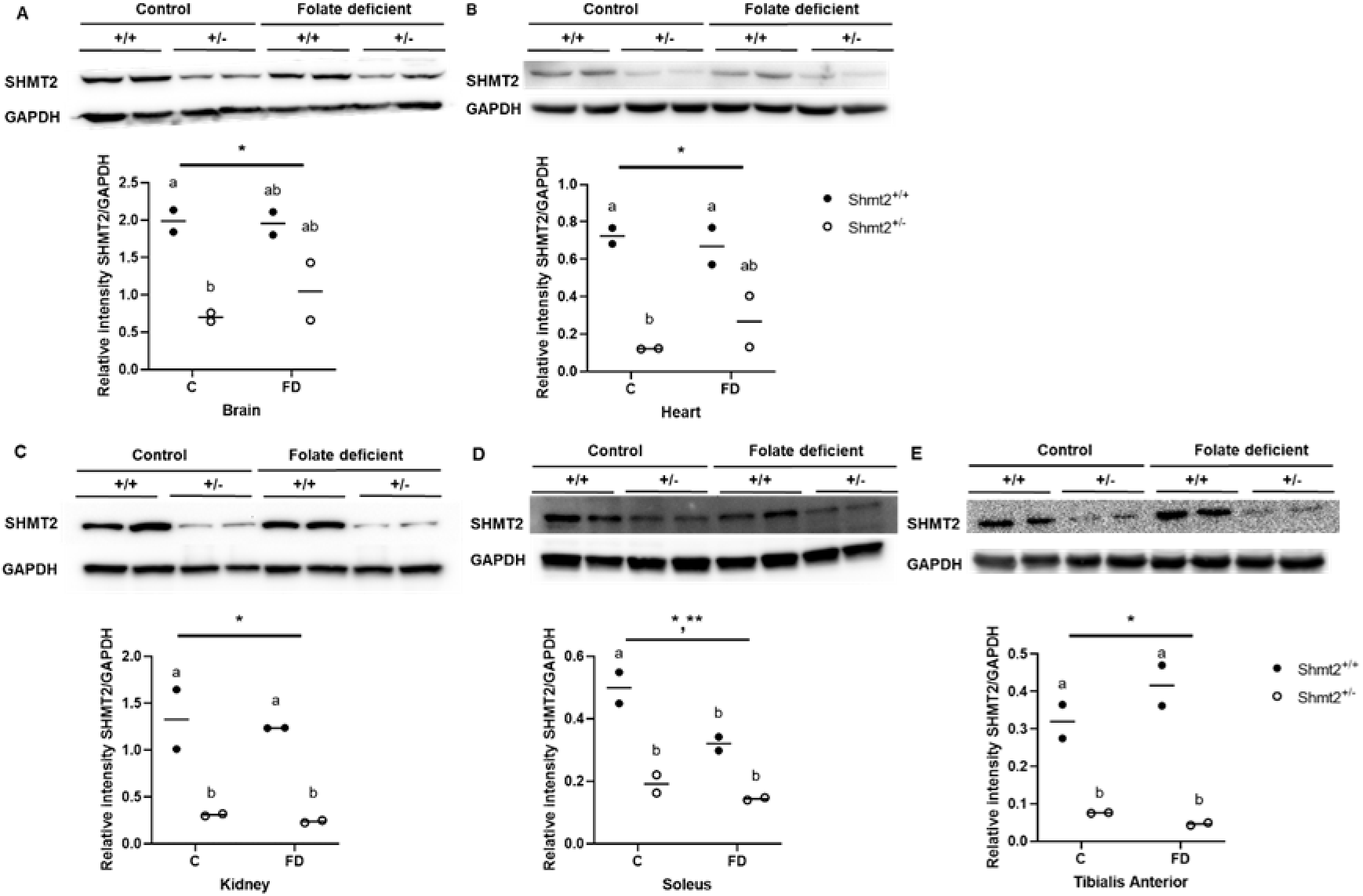
SHMT2 western blot analyses. SHMT2 protein levels in *Shmt2^+/+^* and *Shmt2^+/-^* female mice consuming C or FD diets for 7 weeks in A) brain, B) heart, C) kidney, D) soleus, and E) tibialis anterior. Data from female mice are normalized to GAPDH. Densitometry was performed using ImageJ. Two-way ANOVA with Tukey’s post-hoc analysis was used to determine diet by genotype interaction and main effects of diet and genotype with a statistical significance at *p* ≤ 0.05. Statistical significant was determined for genotype (*), diet (**), and diet by genotype interaction (***). Levels not connected by the same letter are significantly different, n = 2 per group. C, control diet; FD, folate-deficient diet. GAPDH, glyceraldegyde-3 phosphate dehydrogenase; SHMT2, serine hydroxymethyltransferase 2.

### Folate deficiency and *Shmt2^+/-^* genotype influence plasma transmethylation biomarkers but not plasma formate levels

The FD diet significantly increased plasma homocysteine in mice by approximately 50% (**Table 4**). The FD diet was also associated with significantly reduced plasma α-aminobutyric acid, methionine, and dimethylglycine (Table 4), as has been observed previously (19). Additionally, *Shmt2^+/-^* mice exhibited a trend toward slightly higher plasma serine levels (*p* = 0.072). Neither diet nor *Shmt2* genotype influenced plasma levels of cystathionine, cysteine, glycine, or methylglycine. Plasma formate levels were not affected by either the FD diet or *Shmt2^+/-^* genotype though were notably lower than in previous studies (27), likely due to collection of plasma after an overnight fast (Table 4 and **Figure S2**).

**Table 4.** Metabolite profile of plasma from Shmt2^+/+^ and Shmt2^+/-^ mice. Two-way ANOVA with Tukey’s post-hoc analysis was used to determine diet-by-genotype interaction and main effects of diet and genotype with a statistical significance at *p* ≤ 0.05. Data represent means ± SD values. n = 7-10 per group; ns, not significant.

| Metabolite | *Shmt2*<sup>+/+</sup> |  | *Shmt2*<sup>+/-</sup> |  | *p* value of model effect |  |  |
| --- | --- | --- | --- | --- | --- | --- | --- |
|  | C | FD | C | FD | Diet | Genotype | Diet-by-genotype |
| Homocysteine (μM) | 6.6 ± 1.8 | 11.7 ± 3.1 | 5.43 ± 0.9 | 13.0 ± 4.5 | <0.0001 | ns | ns |
| Cystathionine (nM) | 1160 ± 239 | 1162 ± 301 | 1065 ± 238 | 1313 ± 268 | ns | ns | ns |
| Cysteine (μM) | 294 ± 42.0 | 280 ± 27.3 | 277 ± 15.6 | 283 ± 29.0 | ns | ns | ns |
| α-Aminobutyric acid (μM) | 24.2 ± 10.5 | 17.9 ± 2.7 | 30.8 ± 26.9 | 16.4 ± 4.8 | 0.047 | ns | ns |
| Methionine (μM) | 61.4 ± 13.3 | 54.0 ± 3.6 | 61.3 ± 4.7 | 54.9 ± 6.3 | 0.023 | ns | ns |
| Glycine (μM) | 218 ± 15.1 | 231 ± 21.9 | 221 ± 36.8 | 233 ± 27.3 | ns | ns | ns |
| Serine (μM) | 117 ± 12.0 | 108 ± 7.7 | 119 ± 9.6 | 118 ± 12.0 | ns | 0.072 | ns |
| Dimethylglycine (μM) | 13.8 ± 2.2 | 11.2 ± 0.9 | 12.7 ± 2.0 | 11.5 ± 1.5 | 0.006 | ns | ns |
| Methylglycine (μM) | 10.7 ± 4.4 | 12.3 ± 6.0 | 18.3 ± 15.2 | 14.7 ± 5.9 | ns | ns | ns |

### Neither the folate-deficient diet nor decreased SHMT2 levels affect liver mtDNA content or mitochondrial mass

MtDNA copy number and mitochondrial mass were quantified in liver of male mice in response to dietary folate deficiency and decreased SHMT2 levels. No difference in mtDNA content or mitochondrial mass was observed in mouse liver as a result of decreased SHMT2 expression or exposure to the FD diet (**Figure 5A** and 5B).

**Figure 5.**
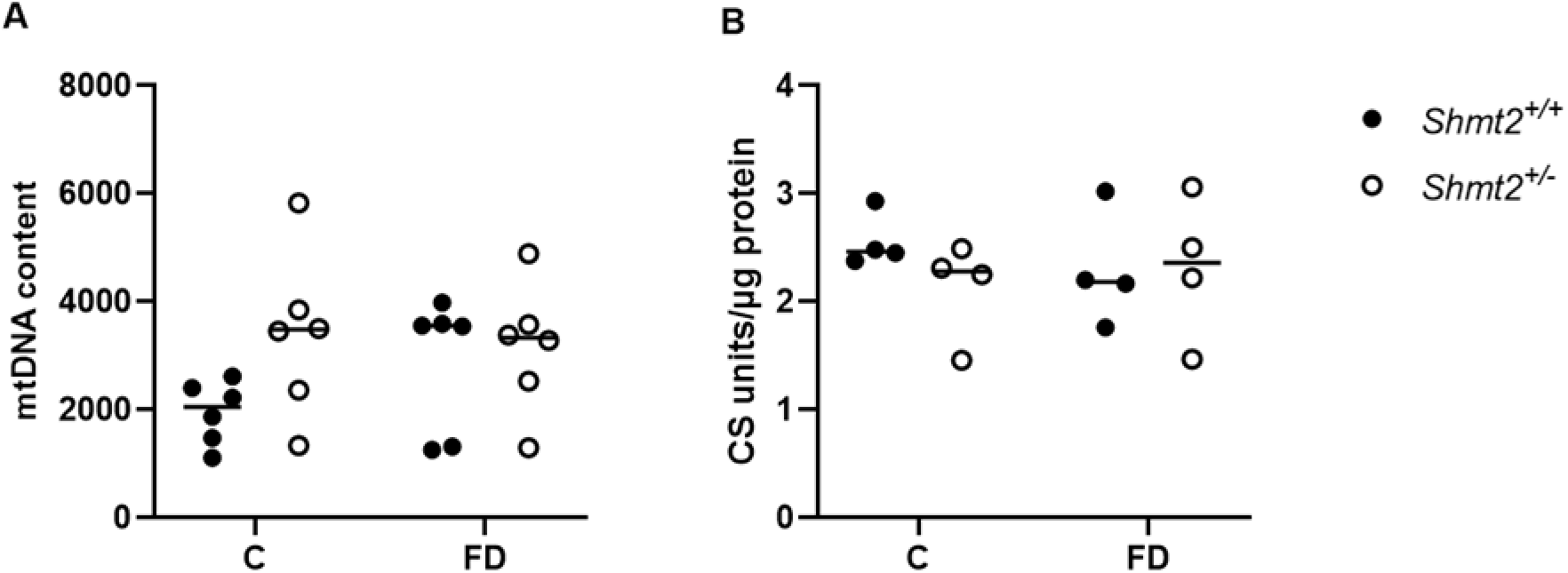
mtDNA content and mitochondrial mass in liver. A) mtDNA content of *Shmt2^+/+^* and *Shmt2^+/-^* mouse liver from male mice consuming the C and FD diets for 7 weeks. Data are normalized using the ΔCT method as previously described (37). Mitochondrial mass determined by citrate synthase (CS) activity assay and normalized to total protein. Two-way ANOVA with Tukey’s post-hoc analysis was used to determine diet by genotype interaction and main effects of diet and genotype with a statistical significance at *p* ≤ 0.05. n = 4-9 per group. There were no significant diet, genotype, or diet-by-genotype interaction in either assay. C, control; FD, folate deficient.

### Uracil levels in liver mtDNA are increased by both reduced *Shmt2* expression and folate deficiency

Limited by the requirement of >2 μg DNA as input, commonly used mass spectrometry-based methods for determining uracil misincorporation into DNA are impractical for use in quantifying uracil in mtDNA. Therefore, we developed a real-time PCR-based assay to assess global uracil content in the mitochondrial genome based on a previously reported method designed for specific loci in nuclear DNA (24). Here, with optimization of real-time PCR conditions, we report a high-sensitivity, long-run real-time PCR technique that enables quantification of total and region specific uracil content in mtDNA (Figure 1B and **Figure S4**).

Total uracil content in mtDNA was significantly increased both by FD diet (*p* < 0.05) and the *Shmt2^+/-^* genotype (*p* < 0.01, **Figure 6A**). There was also an interaction between *Shmt2* heterozygosity and the FD diet (*p* < 0.05, Figure 6A). Of note, rather than evenly distributed throughout the mitochondrial genome (Figure 1B), uracil was mainly incorporated into regions 3-6 (Figure S3) and there was a diet by genotype interaction in region 5 (**Table S1**). In addition, to confirm that the increased total uracil content detected was due to the presence of uracil, mtDNA samples from *Shmt2^+/-^* mice consuming the FD diet were subjected to *in vitro* base excision repair with uracil-DNA glycosylase (UDG) to replace dUTP with dTTP prior to analysis using the novel real-time PCR method. As compared with dTTP repaired mtDNA, untreated mtDNA exhibited significantly higher levels of uracil, validating the specificity of this novel assay (*p* < 0.001, Figure 6B). Importantly, neither the FD diet nor *Shmt2^+/-^* genotype affected uracil content in liver nuclear DNA (**Table 5**).

**Figure 6.**
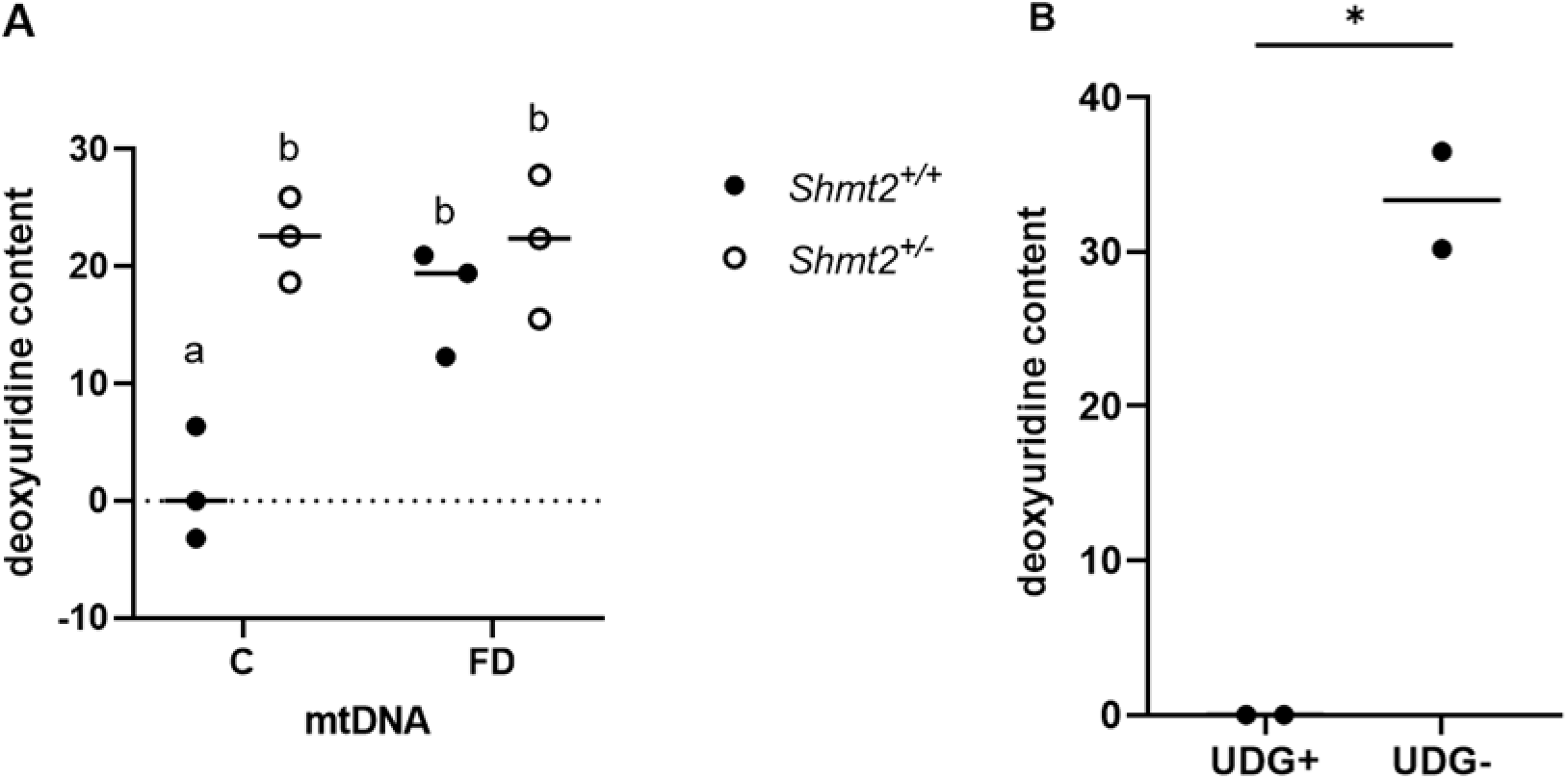
Uracil content in *Shmt2^+/+^* and *Shmt2^+/-^* mouse liver mtDNA. A) Uracil content in *Shmt2^+/+^* and *Shmt2^+/-^* mouse liver mtDNA from male mice consuming the C or FD diet for 7 weeks and B) Uracil content both before and after UDG treatment. The content represents total uracil in each region. Two-way ANOVA with Tukey’s post-hoc analysis was used to determine diet by genotype interaction and main effects of diet and genotype and data are normalized to the *Shmt2^+/+^* mice on control diet in panel A. Statistical significant was determined for genotype (*), diet (**), and diet by genotype interaction (***). Levels not connected by the same letter are significantly different, n = 3 per group. Student’s t-tests were used to analyze data in panel B. Data represent means ± SD. *P* values ≤ 0.05 were considered significantly different. n = 2 per group. * indicates statistical significant. C, control; FD, folate deficient; SHMT2, serine hydroxymethyltransferase 2; UDG, uracil dna glycosylase.

**Table 5.** Liver uracil content in genomic DNA. Two-way ANOVA with Tukey’s post-hoc analysis was used to determine diet-by-genotype interaction and main effects of diet and genotype with a statistical significance at *p* ≤ 0.05. Data represent means ± SD values. n = 4 per group. *There were no significant diet, genotype, or diet-by-genotype interaction in liver uracil content.

| Diet | *Shmt2* genotype | Uracil* |
| --- | --- | --- |
|  |  | *pg uracil/μg DNA* |
| AIN-93G | *Shmt2*<sup>+/+</sup> | 1.05 ± 0.27 |
| | *Shmt2*<sup>+/-</sup> | 1.09 ± 0.31 |
| AIN-93G minus folate | *Shmt2*<sup>+/+</sup> | 1.05 ± 0.42 |
| | *Shmt2*<sup>+/-</sup> | 1.45 ± 0.59 |
| *p* value, diet effect |  | 0.444 |
| *p* value, genotype effect |  | 0.342 |
| *p* value, diet by genotype |  | 0.437 |

### The interaction of *Shmt2* expression and folate availability in mouse embryonic fibroblast (MEF) cells impair cellular proliferation, mitochondrial membrane potential, and mitochondrial function

Examination of protein levels confirmed reduced levels of SHMT2 protein levels were reduced by 65% in *Shmt2^+/-^* MEF cells harvested from (*p* < 0.001, **Figure 7A**). When *Shmt2^+/+^* and *Shmt2^+/-^* MEF cells were grown in modified culture medium containing either 25 nM (6S)5- formylTHF (folate-sufficient medium) or 0 nM(6S) 5-formylTHF (low-folate medium), the accumulation of folate was impaired in cells grown in low-folate medium (*p* < 0.001, Figure 7B). No difference in mtDNA content was observed as a result of either decreased *Shmt2* expression or exposure to the low-folate medium (Figure 7C), consistent with what was observed in liver (Figure 5A). Mitochondrial membrane potential was impaired in the *Shmt2^+/-^* cells and in cells cultured in low-folate culture medium (Figure 7D). Furthermore, *Shmt2^+/-^* cells cultured in either low-folate or folate-sufficient culture medium exhibited reduced proliferative capacity (*p* < 0.05, **Figure 8A**). *Shmt2^+/+^* cells grown in low-folate culture medium also exhibited impaired proliferation (*p* < 0.05, Figure 7A). The addition of formate, but not glycine, rescued impaired proliferation in both *Shmt2^+/+^* cells cultured for three days in low-folate medium and *Shmt2^+/-^* cells cultured in both folate-sufficient or low-folate culture medium (Figure 8B-C). Basal cellular respiration levels (i.e. oxygen consumption rates) were quantified as a biomarker of mitochondrial function. There was a medium-by-genotype interaction in basal respiration levels (*p* < 0.001, **Figure 9A**); *Shmt2^+/+^* MEF cell oxygen consumption was impaired in low-folate culture conditions, and *Shmt2^+/-^* MEF cells exhibited impaired oxygen consumption regardless of folate levels in culture medium. Furthermore, effects of exposure to low-folate medium or decreased *Shmt2* expression on oxygen consumption were not rescued by the addition of glycine or formate (Figure 9A-C).

**Figure 7.**
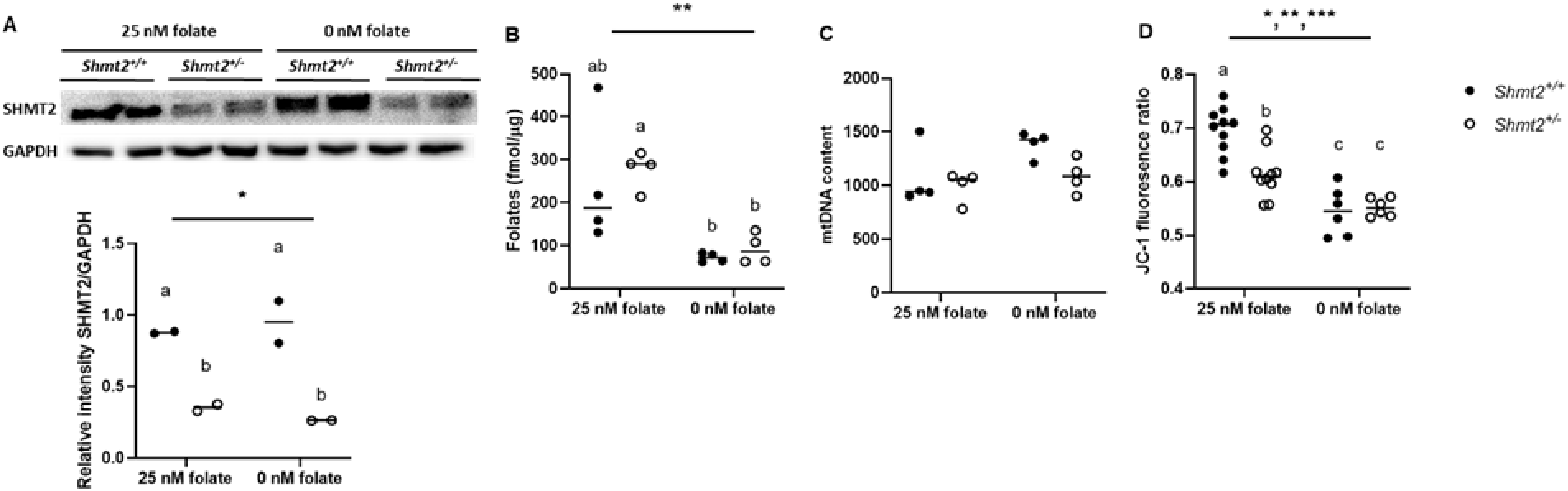
*Shmt2^+/-^* loss and low-folate media decrease mitochondrial membrane potential in MEF cells. A) SHMT2 protein levels, B) total folate levels, C) mtDNA content, and D) mitochondrial membrane potential. SHMT2 protein levels were normalized to GAPDH and densitometry was performed using ImageJ. Two-way ANOVA with Tukey’s post-hoc analysis was used to determine media by genotype interaction and main effects of media and genotype with a statistical significance at *p* ≤ 0.05. Statistical significance was determined for genotype (*), diet (**), and diet by genotype interaction (***). Levels not connected by the same letter are significantly different, n = 2-8 per group. GAPDH, glyceraldegyde-3 phosphate dehydrogenase; SHMT2, serine hydroxymethyltransferase 2.

**Figure 8.**
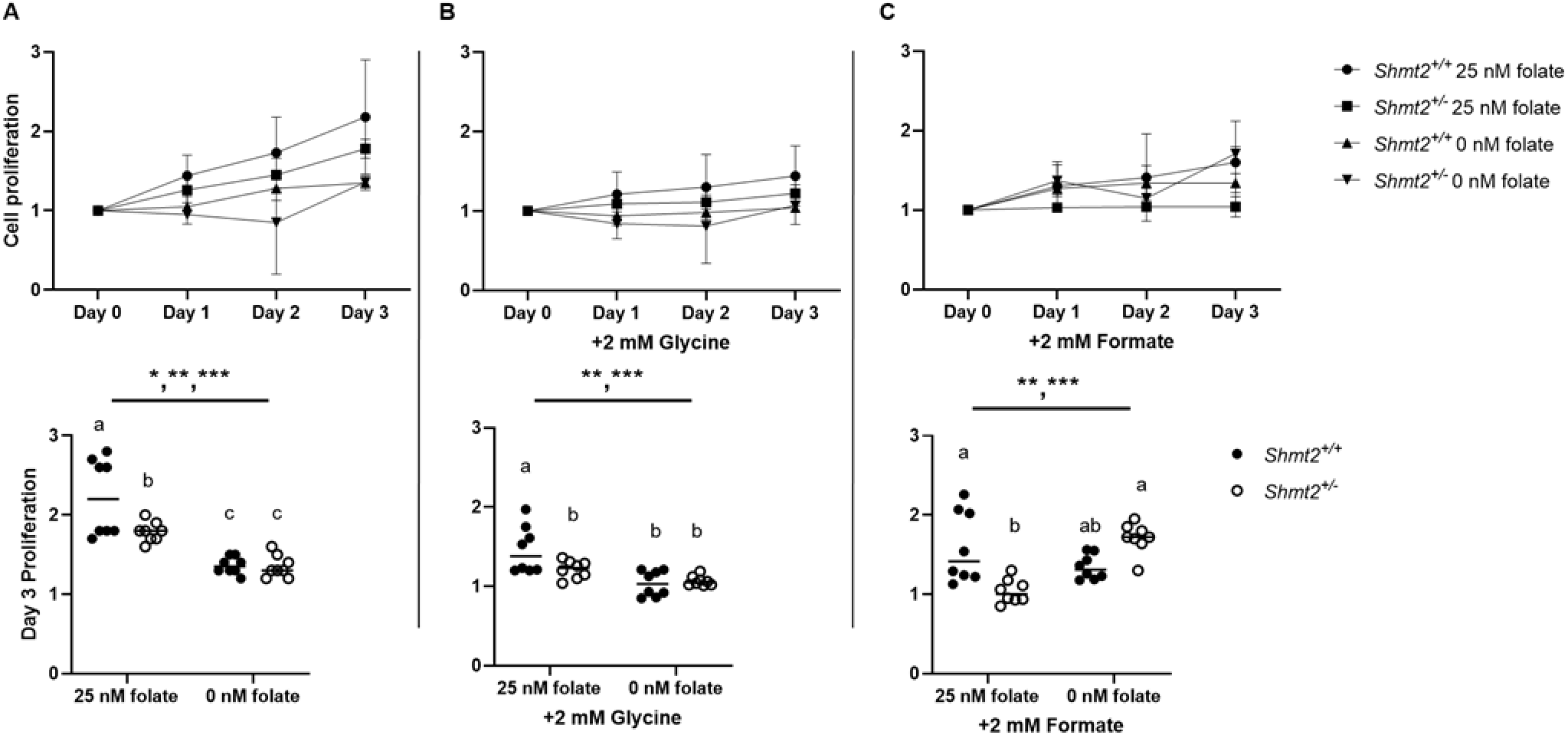
*Shmt2^+/-^* loss and low-folate culture medium decrease cell proliferation rate in MEF cells. Cell proliferation rates of *Shmt2^+/-^* MEF cells were compared with *Shmt2^+/+^* MEF cells by co- staining cells with Hoechst 33342 (to identify all cells) and propidium iodide (to identify dead cells). Fold change of each group was calculated by dividing by day 0 cell number. Values represent n=4 replicates of 2 embryo cell lines cultured in medium containing either 25 nM (6S)5-formylTHF or 0 nM (6S)5-formylTHF. A) Cell proliferation rate and day 3 quantitation, cell proliferation rate in the presence of 2 mM glycine and day 3 quantitation, and C) cell proliferation in the presence of 2 mM formate and corresponding day 3 quantitation. Linear mixed effects models with random effect of embryo, main effects of media, genotype, and time (with time as a continuous variable), and all 2-, 3-, and 4-way interactions were used to determine cell proliferation with a statistical significance at *p* ≤ 0.05. Two-way ANOVA with Tukey’s post-hoc analysis was used to determine media by genotype interaction and main effects of media and genotype with a statistical significance at *p* ≤ 0.05 were used to analyze day 3 proliferation. Statistical significance was determined for genotype (*), diet (**), and diet by genotype interaction (***). Levels not connected by the same letter are significantly different.

**Figure 9.**
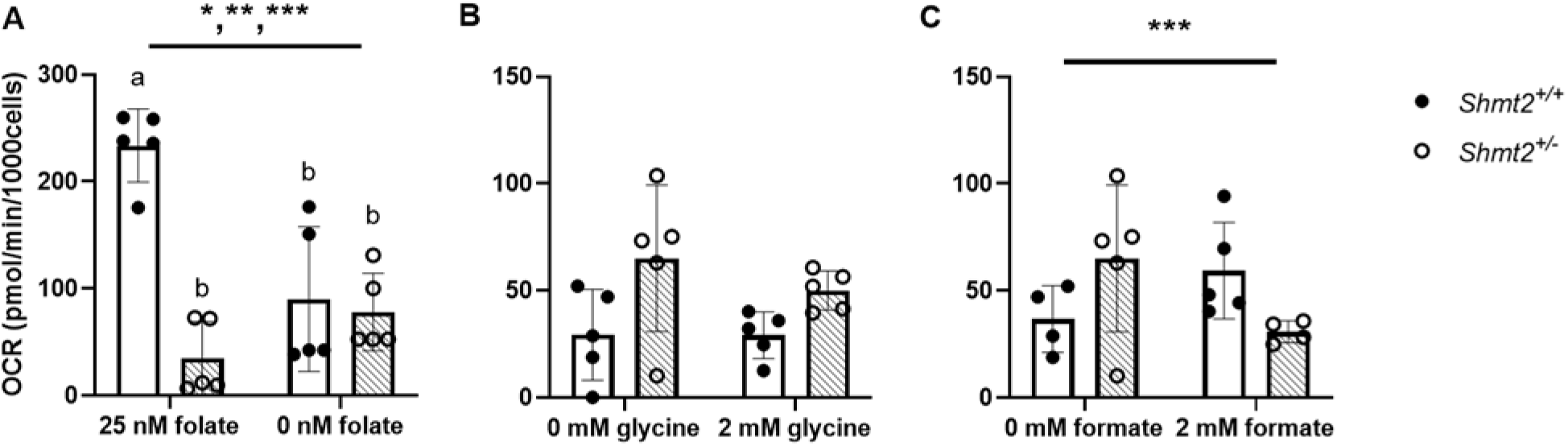
*Shmt2^+/-^* loss and low-folate media decrease mitochondrial oxygen consumption rate in MEF cells. A) oxygen consumption rate (OCR) of *Shmt2^+/+^* and *Shmt2^+/-^* MEF cells were cultured in medium containing either 25 nM (6S)5-formylTHF or 0 nM (6S)5-formylTHF, B) oxygen consumption rate of *Shmt2^+/+^* and *Shmt2^+/-^* MEF cells grown in culture medium containing 0 nM (6S)5-formylTHF and 0 or 2 mM glycine for 48 hours, and C) oxygen consumption rate of *Shmt2^+/+^* and *Shmt2^+/-^* MEF cells grown in culture medium containing 0 nM (6S)5-formylTHF and 0 or 2 mM formate for 48 hours. Two-way ANOVA with Tukey’s post-hoc analysis was used to determine media by genotype interaction and main effects of media and genotype with a statistical significance at *p* ≤ 0.05. Statistical significance was determined for genotype (*), diet (**), and diet by genotype interaction (***). Levels not connected by the same letter are significantly different, n = 4-5 per group.

## DISCUSSION

In this study, a whole-body *Shmt2* knockout mouse was generated to study the effects of reduced SHMT2 protein/activity levels on mitochondrial DNA integrity, mitochondrial function, and energy metabolism. Consistent with observations by Tani *et. al.*, homozygous *Shmt2* knockouts are embryonic lethal, demonstrating the essentiality of *Shmt2* (Table 2) (26). Biallelic human variants in *SHMT2* have recently been identified in a small number of patients using whole-exome sequencing (28). Most of these variants are hypothesized to affect substrate binding and/or SHMT2 protein oligomerization and ultimately protein function (28). Biallelic SHMT2 variants result in neurological complications, among other phenotypes, in affected individuals. Some of these variants are commonly inherited single-nucleotide polymorphisms. However, the effects of these variants inherited as single alleles, which are likely to more subtly impair SHMT2 function, on mitochondrial capacity and tissue function remain unclear. *Shmt2^+/-^* mice exhibit decreased SHMT2 protein levels across a range of tissues (Figures 3-4), and therefore provide a model of these more modest SHMT2 impairments.

Total folate accumulation in the plasma was reduced in mice consuming a FD diet (Table 3). There was also a significant interaction of diet and genotype suggesting the reduced folate levels were more pronounced in the *Shmt2^+/+^* mice consuming FD diet (88%) compared to the *Shmt2^+/-^* mice (80%). Unexpectedly, *Shmt2^+/-^* mice consuming sufficient dietary folate had reduced total folate levels in liver mitochondria. This finding suggests that decreased SHMT2 levels impair mitochondrial folate accumulation, though the underlying mechanisms remain uncharacterized. Interestingly, *Shmt2^+/-^* mice also exhibit increased uracil in liver mtDNA even when consuming adequate dietary folate, and dietary folate deficiency increases uracil content in mouse liver mtDNA in both *Shmt2^+/+^* and *Shmt2^+/-^* mice (Figures 6 and S3), without impacting uracil in liver nuclear DNA. The uracil misincorporation in mtDNA was in alignment with previous findings in transformed/immortalized cell models demonstrating that cells with reduced SHMT2 protein levels or cultured in the presence of low-folate culture medium exhibit increased uracil in mtDNA (10, 16). Furthermore, uracil does not appear evenly distributed throughout the entire mitochondrial genome, but appears enriched within the certain regions of the mtDNA (Figure S3), highlighting the need to develop high-resolution methods to interrogate the distribution patterns of uracil in mtDNA. Interestingly, the effects of reduced *Shmt2* expression and folate deficiency do not seem to be additive with respect to uracil accumulation in mtDNA. This observation suggests that liver mtDNA can tolerate uracil accumulation to a certain threshold, though mechanisms governing this threshold (and whether this threshold applies to all tissues) remain unidentified. One of the unique features of mtDNA is that it is polyploid, with up to several thousand copies of their genome per cell (29). The high copy number of mtDNA provides genetic redundancy that may “buffer” mtDNA mutations/lesions to maintain mitochondrial function. Therefore, it is possible that a given “threshold” level of uracil misincorporation is necessary before biochemical defects or mitochondrial dysfunction becomes apparent. Consistent with the idea that the susceptibility of cells to developing mitochondrial dysfunction is influenced by the copy number of mtDNA, it has been shown that the synthesis and degradation of mtDNA is highly regulated to maintain mtDNA copy number and the turnover rate varies depending on the tissues. Rat heart, liver, and kidney exhibit the highest mtDNA turnover rates, whereas brain has a markedly lower mtDNA turnover rate (29, 30). Thus, it is possible that the mtDNA content in liver is maintained by rapid mtDNA turnover while uracil is continuously misincorporated into mtDNA during this process. Although this study did not examine uracil levels in tissues with low mtDNA turnover rates, it is possible decreased SHMT2 could lead to more pronounced affects in those more energetic tissues.

Decreased *Shmt2* expression in immortalized cell models leads to inconsistent changes in mtDNA content with some models exhibiting increased mtDNA content in response to decreased *SHMT2* expression (16) and some models reporting no response in mtDNA content (31, 32). Our results suggest mtDNA is sensitive to uracil misincorporation in response to impaired mitochondrial folate metabolism, whereas mtDNA content may be modulated in a tissue/cell-type specific manner.

Reduced *Shmt2* expression and exposure to low-folate medium in MEF cells significantly impaired cellular respiration, likely through multiple mechanisms and ultimately reduced proliferation rates. These impairments included both reduced mitochondrial membrane potential and oxidative capacity (Figures 7D and 9A). This data is consistent with other MEF and immortalized cells models of homozygous *SHMT2* loss or folate deficiency that exhibit decreased oxidative capacity and impaired mitochondrial complex I activity and protein levels, suggesting FOCM and the oxidative phosphorylation system are functionally coordinated (26,31–34). Furthermore, the reduced membrane potential is consistent with the observations in human cells with biallelic *SHMT2* variants (28). In immortalized cell models, SHMT2-induced energy metabolism changes are thought to be a result of impaired mitochondrial translation (32), reduced complex I protein levels (26, 31), or decreased NADPH production (34). Lucas *et. al.* not only found *SHMT2* loss impairs complex I protein levels in 293A cells, but also found the addition of formate, but not glycine, was able to rescue this impairment (31). In MEF cells decreased *Shmt2* expression and exposure to low-folate medium impaired proliferation and oxygen consumption rates. Glycine failed to restore either cell proliferation or mitochondrial function (Figure 8E and 9B). In contrast, formate restored cellular proliferation (Figure 8C) in *Shmt2^+/+^* MEF cells maintained in the low-folate medium but did not affect proliferation of *Shmt2^+/-^* MEF cells. Additionally, formate was not able to restore oxygen consumption rates in cells cultured in low-folate medium, regardless of *Shmt2* genotype (Figure 9A and 9C). Taken together, these data indicate that provision of one-carbon units is insufficient to restore mitochondrial function as a result of impaired mitochondrial function in MEF cells. These findings also highlight key differences in response to impaired mitochondrial folate metabolism between transformed/immortalized and primary cells.

One limitation of this study is that although the CRISPR-Cas9 targeting strategy used to generate *Shmt2^+/-^* mice likely reduced expression of all isoforms of SHMT2 (i.e. the pre- processed form and SHMT2α isoform), currently available antibodies detected only the processed, mitochondrial SHMT2. Since SHMT2α contributes to cytosolic/nuclear dTMP synthesis (35), it is possible that reduced SHMT2α levels are also modulating total cellular dTMP synthesis. Though it is worth noting that any reduction in SHMT2α in this model did not affect uracil levels in nuclear DNA (Table 5). Another limitation of this study is that the effects of loss of *Shmt2* on mtDNA integrity have only been assessed in the mouse liver. Given that differential responses to loss of SHMT2 were observed from different human cancer cell lines (26,31–34), it is plausible that SHMT2 contributes to mtDNA maintenance in a tissue-specific manner, warranting further investigation of other mouse tissues, particularly tissues that rely heavily on mitochondrial function or have low mtDNA turnover rates, such as brain, heart, and skeletal muscle. In addition, since age-related changes in mtDNA are associated with mitochondrial function decline and SHMT2 levels are approximately 50% lower in aged fibroblasts (36), the effects of reduced *Shmt2* expression on mtDNA integrity in aged mice should also be assessed in future work. In summary, we have demonstrated that total uracil content of mouse liver mtDNA is increased as a result of both *Shmt2* heterozygosity and dietary folate deficiency. In addition, *Shmt2* heterozygosity and exposure to low-folate culture medium affect mitochondrial function and proliferation in MEF cells.

## Acknowledgements

We acknowledge and thank Lynn Johnson at the Cornell Statistical Consulting Unit for her help with the statistical analyses for this manuscript. We would also like to acknowledge Kaydine Edwards of the Cornell University Division of Nutritional Sciences for assistance with Agilent Seahorse XF Analyzer experiments and cell culture, respectfully.

## Author contributions

JLF, YX, and MSF designed research; JLF, YX, JEB, SGL, WEP, SPS, MEB, JTB, ATM conducted research; JLF, YX, and MSF analyzed data; JLF and MSF prepared the manuscript; and MSF has primary responsibility for the final content. All authors read and approved the final manuscript.

