## Supplementary Information for "Reduced *Shmt2* expression impairs mitochondrial folate accumulation and respiration, and leads to uracil accumulation in mouse mitochondrial DNA"

Table S1

***P* values for deoxyuridine content in *Shmt2^+/+^* and *Shmt2^+/-^* mouse liver mitochondrial DNA.** Two-way ANOVA with Tukey’s post-hoc analysis was used to determine diet-by-genotype interaction and main effects of diet and genotype with a statistical significance at *p* ≤ 0.05. Total U is the uracil *p* value for the total mtDNA while regions 1 through 6 represent specific mtDNA regions (as described in the methods).

|  | Total U | Region 1 | Region 2 | Region 3 | Region 4 | Region 5 | Region 6 |
| --- | --- | --- | --- | --- | --- | --- | --- |
| Diet | 0.036 | 0.050 | 0.039 | 0.570 | 0.020 | 0.870 | 0.285 |
| Genotype | 0.002 | 0.031 | 0.230 | 0.590 | 0.069 | 0.250 | 0.511 |
| Diet-by-genotype | 0.040 | 0.065 | 0.466 | 0.376 | 0.063 | 0.030 | 0.187 |

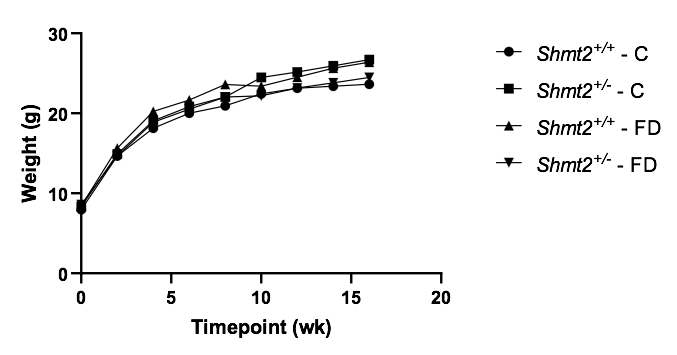

Supplemental Figure 1. Growth curves. Growth curves analysis for *Shmt2^+/+^* and *Shmt2^+/-^* male and female mice from weaning to adulthood. At weaning, mice were placed on the C or FD diet and body weights were measured at weaning and every two weeks thereafter. A linear mixed model was used to determine the fixed effects of genotype, diet, and the interaction between genotype and diet. The model allowed for separate intercepts, slopes, and quadratic terms of time for each genotype by diet group and random intercepts, slopes, and quadratic effects at the mouse level. A residual analysis was performed to check the model assumptions of normality and homogeneous variance. F tests using the Kenward-Roger degrees of freedom correction were used to test the statistical significance of the model fixed effects. Data represent means at each time point. *P* values ≤ 0.05 were considered significantly different. n = 3-7 per group. C, control diet; FD, folate-deficient diet.

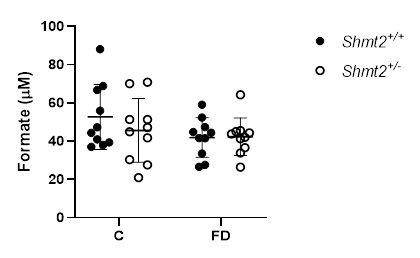

Supplemental **Figure 2** **Formate content in *Shmt2^+/+^* and *Shmt2^-/+^* female mouse plasma.** Two-way ANOVA with Tukey’s post-hoc analysis was used to determine diet-by-genotype interaction and main effects of diet and genotype. There were no significant differences (*p* > 0.05), n = 10 per group.

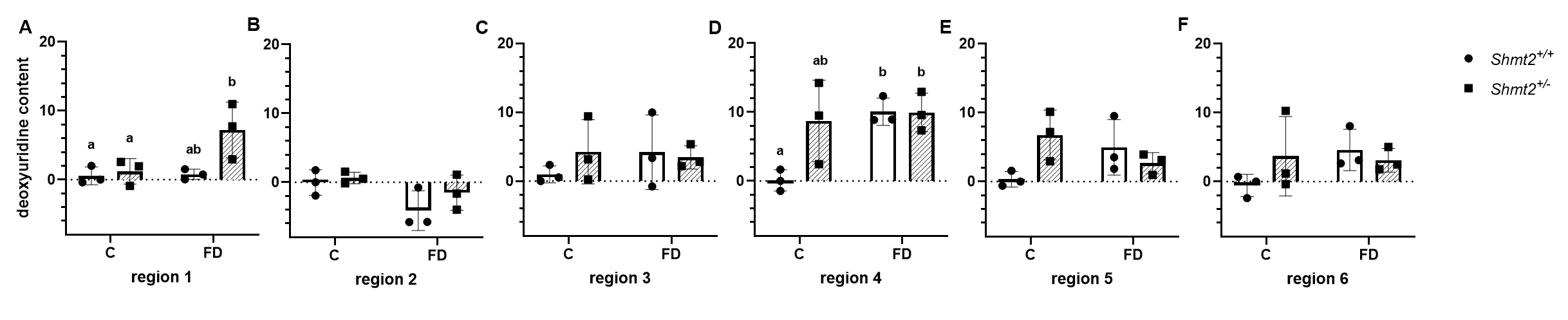

Supplemental **Figure 3 Uracil content in *Shmt2^+/+^* and *Shmt2^-/+^* mouse liver mitochondrial DNA regions 1 through 6.** Two-way ANOVA with Tukey’s post-hoc analysis was used to determine diet by genotype interaction and main effects of diet and genotype with a statistical significance at *p* ≤ 0.05. Statistical significant was determined for genotype, diet, and diet-by-genotype interaction within each region and reported in Table S1. Levels not connected by the same letter are significantly different, n = 3 per group.

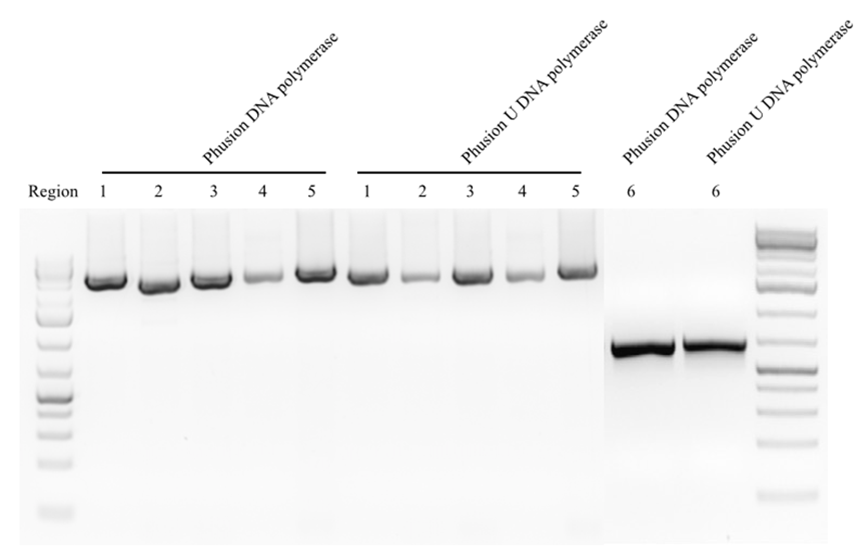

Supplemental **Figure 4 PCR products in *Shmt2^+/+^* and *Shmt2^+/-^* mouse liver mtDNA.**

Test of specificity of long-run real-time PCR. All primers from Table 1 yielded single products in both *Shmt2^+/+^* and *Shmt2^+/-^* mouse liver mtDNA.
